# Cell competition, the kinetics of thymopoiesis and thymus cellularity are regulated by double negative 2 to 3 early thymocytes

**DOI:** 10.1101/2019.12.21.885673

**Authors:** Camila V. Ramos, Luna Ballesteros-Arias, Joana G. Silva, Rafael A. Paiva, Marta F. Nogueira, Jorge Carneiro, Erida Gjini, Vera C. Martins

## Abstract

Cell competition in the thymus is a homeostatic process that drives turnover. If the process is impaired, thymopoiesis can be autonomously maintained for several weeks, but this causes leukemia. We aimed to understand the impact of cell competition on thymopoiesis, identify the cells involved and determine how the process is regulated. Using thymus transplantation experiments we found that cell competition occurs within the double negative 2 (DN2) and 3 early (DN3e) thymocytes and inhibits thymus autonomy. Furthermore, the expansion of DN2b is regulated by a negative feedback loop imposed by double positive thymocytes and determines the kinetics of thymopoiesis. This feedback loop impacts on cell cycle duration of DN2b, in a response controlled by interleukin 7 availability. Altogether, we show that thymocytes do not merely follow a pre-determined path if provided with the correct signals. Instead, thymopoiesis dynamically integrates cell autonomous and non-cell autonomous aspects that fine-tune normal thymus function.

## INTRODUCTION

The thymus is an organ with high cellular turnover in which virtually all thymocytes are replaced every four weeks (Berzins et al., 1998). Progenitors of bone marrow origin enter the thymus at the cortico-medullary junction, commit to the T cell lineage, and progressively go through a series of differentiation stages before becoming T lymphocytes (Yui and Rothenberg, 2014). Each stage is brief and thymocytes are constantly replaced by new incoming cells (Petrie and Zuniga-Pflucker, 2007). Differentiation stages can be distinguished with basis on the expression of surface markers, with the major subpopulations defined by differential expression of CD4 and CD8 co-receptors. Double negative (DN) thymocytes lack both CD4 and CD8. As they mature, CD8 is upregulated to generate immature single positive (ISP) thymocytes. CD4 is expressed afterwards to originate CD4+CD8+ double positive thymocytes, which then downregulate one of the co-receptors as they turn into CD4, or CD8 single positive thymocytes, that subsequently leave the thymus. The DN stages can be further detailed and the most immature thymocytes are early T lineage progenitors (ETP), defined by high expression of Kit/CD117 and CD44. These cells upregulate CD25 as they differentiate into DN2a and then DN2b, which express high or intermediate kit/CD117, respectively. DN2 progressively loose CD117 and CD44 expression when they differentiate into DN3. The progression from DN3 to DN4 is linked with the loss of CD25 expression, which then differentiate into ISP as stated before (Yui and Rothenberg, 2014). T lymphocyte differentiation is associated with thymocyte migration from the cortico-medullary junction through the cortex towards the sub-capsular zone, then back through the cortex, and finally into the medulla (Lind et al., 2001; Porritt et al., 2003). Migration in the thymus is regulated and reflects the existence of several niches, each of them supplying distinct signals that are required for the stage-specific differentiation of thymocytes (Petrie and Zuniga-Pflucker, 2007; Takahama et al., 2017).

Normal thymopoiesis relies on a continuous seeding of the thymus by uncommitted progenitors of bone marrow origin (Love and Bhandoola, 2011; Rothenberg, 2019). Notwithstanding, thymus transplantation experiments revealed that if deprived of *de novo* progenitor seeding, the thymus is capable of maintaining T lymphocyte production independently of bone marrow contribution for several weeks (Martins et al., 2012; Peaudecerf et al., 2012), a condition that we termed thymus autonomy. In the steady state, thymus autonomy is actively inhibited by cell competition. Specifically, thymocytes at the same differentiation stage but with shorter time of thymus residency outcompete and replace those with longer dwell time in the thymus, thereby promoting turnover (Martins et al., 2014). Failure in cell competition can be achieved experimentally in thymi grafted into recipients that either lack progenitors capable of thymic colonization, or that are deficient for interleukin 7 receptor (IL-7r) (Ballesteros-Arias et al., 2019; Martins et al., 2014). In either condition, the thymus grafts have a period of thymus autonomy that is followed by T cell acute lymphoblastic leukemia (T-ALL) with onset at 15 weeks post transplantation and incidence reaching up to 80% (Ballesteros-Arias et al., 2019; Martins et al., 2014). Such results illustrate the importance of cell competition in the maintenance of thymus homeostasis.

Cell competition was originally described in *Drosophila* (Morata and Ripoll, 1975) as a cellular interaction contributing to tissue homeostasis and optimal organ function (Baker, 2017; Johnston, 2009; Morata and Ballesteros-Arias, 2014; Moreno, 2008). It considers that if cellular heterogeneity occurs in a tissue, the least fit cells are eliminated while the fittest prevail and contribute in full for the tissue. Loser cells are viable and capable of maintaining tissue function. Their elimination results from the presence of the fittest cells in close proximity in the same tissue. Cell competition has been further validated in organ homeostasis in rats and mice (Claveria and Torres, 2016; Di Gregorio et al., 2016; Maruyama and Fujita, 2017). In the hematopoietic system, cell competition was found to maintain a fit population of hematopoietic stem cells following stress induced by mild irradiation (Bondar and Medzhitov, 2010). While it is clear that cell competition is an important homeostatic process, there is evidence for several molecular mechanisms depending on the cellular context (Claveria and Torres, 2016; Maruyama and Fujita, 2017).

Here we sought to address the conditions that regulate thymus turnover and study how thymocyte subpopulations maintain stable thymopoiesis. Using thymus transplantation experiments we show that cell competition takes place in the thymic cortex, involves the double negative 2 (DN2) and 3 early (DN3e) thymocytes and results in the inhibition of thymus autonomy. Response to IL-7 defined the cellularity of these populations, suggesting that cell numbers play a role in the functional outcome of cell competition. In addition, we show that DN2b act as homeostatic sensors by transiently adjusting cell numbers in response to a negative feedback loop imposed by double positive thymocytes, in a response that was regulated by IL-7 availability. This response of the DN2b defined the kinetics of thymus turnover. Specifically, DN2b transiently adjusted absolute cell numbers to the time requirement for the differentiation of double positive thymocytes. This impacted progressively on the cell numbers of the following differentiation stages, slowing down thymopoiesis, when double positive thymocytes remained longer at that stage. By comparing thymopoiesis between identical hosts, and changing the genotype of thymus donors, we could determine that thymocytes do not merely follow a pre-determined path if provided with the correct signals. Instead, thymopoiesis results from the integration of cell autonomous and non-cell autonomous aspects, both contributing to the kinetics of differentiation of T lymphocytes. DN2 to DN3e stages are central in adjusting thymopoiesis while also mediating cell competition, both integrated to ensure that overall cellularity does not change dramatically.

## RESULTS

### Thymus turnover is regulated by cell competition in double negative thymocytes

Upon deprivation of fit bone marrow derived progenitors, the thymus is capable of autonomously maintaining T lymphocyte production but T-ALL emerges as a consequence (Ballesteros-Arias et al., 2019; Martins et al., 2014). To identify the cell populations that are involved in cell competition in the thymus, we performed thymus transplantation experiments (Fig. 1A). Upon transplantation of wild type thymus grafts into wild type hosts, the most immature populations of thymocytes, i.e. ETP and DN2a, both defined by high expression of CD117, differentiated and by seven days after transplantation no ETP and DN2a of thymus donor origin could be detected (Fig. 1B). At that time point, no thymocytes of host origin could be detected in the thymus grafts. One week later, by 14 days post-transplantation, ETP and DN2a populations were completely composed of host derived cells, indicating that the differentiation of these populations progresses normally upon transplantation and that donor and host ETP and DN2a do not coexist (Fig. 1B). This observation was consistent in all conditions tested, regardless of the genotype of the donors and of the hosts (not shown). From the DN2b (CD117-intermediate) stage of differentiation onwards, we could observe that donor and host derived thymocytes coexisted, and detail the kinetics with which host derived cells progressively replaced donor thymocytes (Fig. 1C, D). To define the spatial localization of thymocytes, we analyzed thymus grafts at various time points by immunohistology (Fig. 1E). DN2 and DN3 cells were detected by CD25 expression and co-staining with CD45.2 identified the host thymocytes (Fig. 1E). Cortical and medullary areas were distinguished by cytokeratin 8 (K8) staining that labels cortical thymic epithelial cells (Fig. 1E). Thymocytes followed the normal, well-known migratory trajectory throughout the thymus (Fig.1E). Specifically, CD25-positive thymocytes, i.e. DN2 and DN3, were dispersed in the cortex and accumulated at the sub-capsular zone (Fig. 1E). Overall results for the kinetics of turnover were similar to those obtained by flow cytometry (Fig. 1F). The quantification of the thymus sections showed that at stages when both donor and host derived CD25-positive thymocytes could be detected, their spatial distribution was consistent with the migration of donor derived cells, with longer time of thymus residency, ‘ahead’ of the host derived cells. Hence, host thymocytes were detected at the sub-capsular zone at later time points of analysis (Fig. 1G). Altogether, these data show that ETP and DN2a differentiate before there is *de novo* thymus seeding and is therefore unlikely that these donor populations could persist in the thymus in a setting of progenitor deprivation. Donor- and host-derived thymocytes coexist in time starting at DN2b, but not earlier stages. Furthermore, donor and host DN2b and DN3 can locate to the same regions of the thymus, making it plausible that they interact directly, and that such interaction inhibits autonomy.

**Figure 1.**
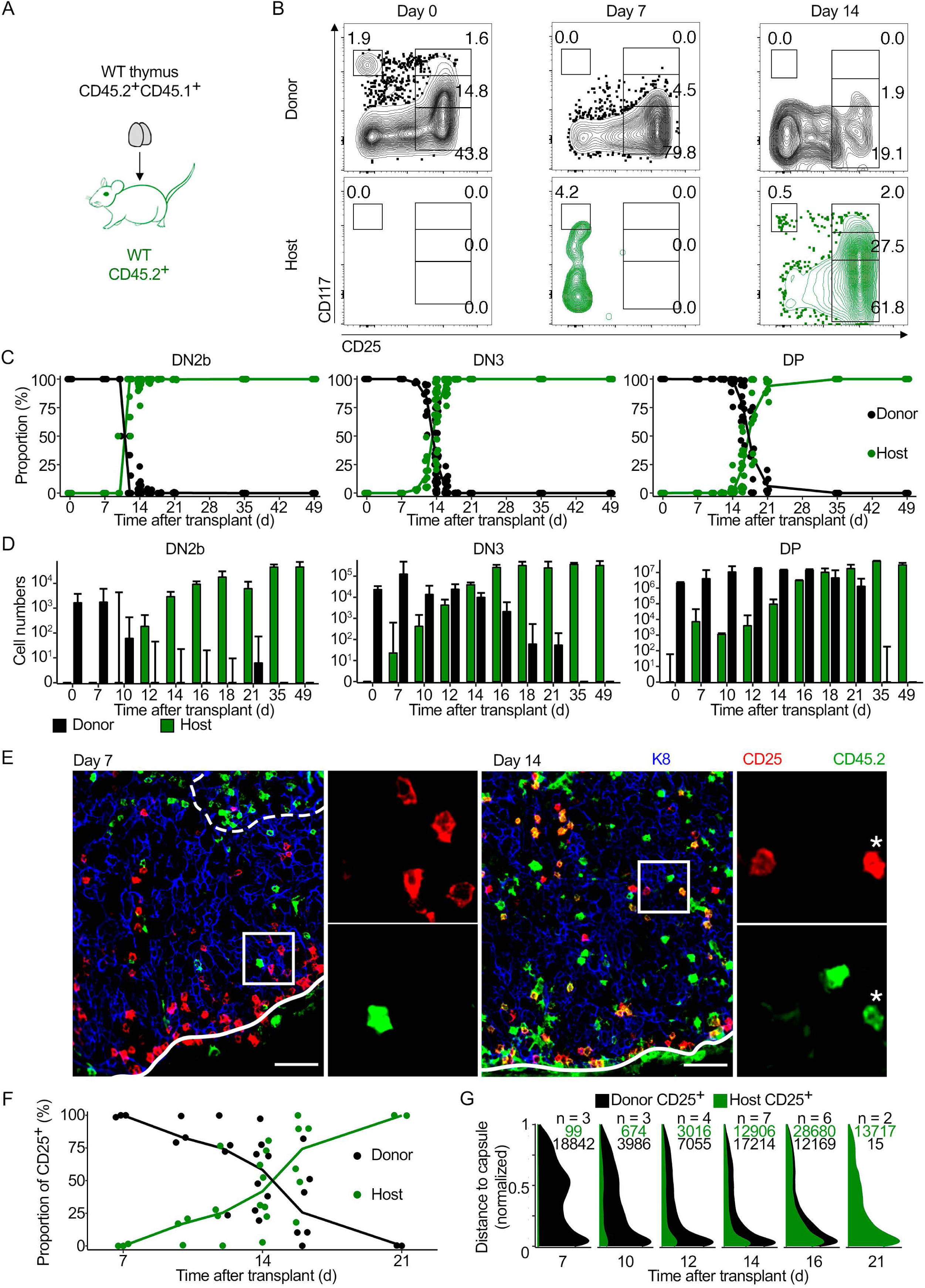
Kinetics of thymus turnover in thymus transplants. **(A)** Schematics of the experimental design. Wild type (WT) newborn thymi were transplanted under the kidney capsule of adult wild type (WT) recipients. **(B)** Thymus grafts were analyzed by flow cytometry at 1-week intervals. Depicted is the expression of CD117 and CD25 in donor (CD45.2+ CD45.1+) or host (CD45.2+) thymocytes pre-gated for the markers CD4^-^CD8^-^Lineage^-^. Gated populations from top left to bottom right are ETP, DN2a, DN2b, and DN3. **(C)** Quantification of the percentage of donor (black circles and line) and host (green circles and line) thymocytes in the indicated populations (DP = CD4+CD8+ double positive). Every thymus graft is represented by a symmetric pair of (black and green) circles. **(D)** Absolute number of DN2b (left), DN3 (middle) and CD4+CD8+ double positive (DP, right) were determined at the indicated time points after thymus transplant (in days, d). Donor and host derived thymocytes are represented in black or green bars, respectively. Data were pooled from several experiments using 6-8 grafts in total for most time points and up to 24 grafts at day 14. Shown is the median +95% confidence interval. **(E)** Thymus grafts were analyzed by immunohistology for the indicated surface markers, CD25 (red), CD45.2 (green) and cytokeratin 8 (K8, in blue). Depicted is one representative example analyzed 7 (left) or 14 days (right) after transplant. Magnification of 20X, scale bars = 50μm. Experiments analyzed by immunohistology followed the experimental design in (A) but donors were B6/Ly5.1 (CD45.1). Asterisk indicates a CD25+ cell of host origin (CD25 positive CD45.2 positive, in yellow). **(F)** Quantification of the relative proportion of donor (black circles and line) and host (green circles and line) cells in CD25+ thymocytes from thymus sections at different time points post-transplant. Each pair of circles (green and black) represents one thymus graft. **(G)** Stacked density distribution of the distance of donor and host-derived CD25+ cells to the capsule (normalized by the maximum distance observed) at the indicated time points. Host CD25+ are depicted in green and donor CD25+ in black. N is the number of grafts analyzed, and numbers correspond to the number of CD25 positive thymocytes of donor (black) or host (green) origin at the indicated time points.

### Cell competition involves thymocytes up to the DN3 early

Classical thymus transplantation experiments into immunodeficient hosts failed to reveal thymus autonomy because the host hematopoietic progenitors could colonize the thymus grafts and initiate thymopoiesis (Frey et al., 1992; Takeda et al., 1996). We reasoned that such experimental setting could be informative about the thymocyte populations involved in cell competition and how they impact on thymopoiesis. To address that, we grafted wild type donor thymi under the kidney capsule of *Rag2*^*-/-*^ recipients (Fig.2A). *Rag2*^*-/-*^ thymocytes cannot rearrange their T cell receptor *loci* and therefore fail to differentiate beyond the DN3 early (DN3e) stage. Similar to the previous experiments, we never detected coexistence of donor and host thymocytes at the ETP and DN2a stages (not shown). On the other hand, DN2b and DN3e thymocytes were detected in coexistence at various time-points after transplantation, and the differentiation and emigration of wild type T lymphocytes from the thymus grafts in *Rag*-deficient recipients occurred with kinetics that were similar to those observed in wild type hosts (Fig. 1D, 2B). The only difference was that thymopoiesis in *Rag2*^*-/-*^ recipients ceased after one single wave of donor T lymphocyte production, as expected, since *Rag*-deficient thymocytes are developmentally arrested at the DN3e stage. Since donor cells do not persist beyond the DN3 stage, the inhibition of thymus autonomy by cell competition must occur before, i.e. between thymocytes up to the DN3e stages of differentiation. Furthermore, the fact that the analyzed wave of thymopoiesis was similar in wild type and in *Rag2*^*-/-*^ recipients suggests that T lymphocyte differentiation downstream of the DN3 is mostly regulated cell autonomously.

**Figure 2.**
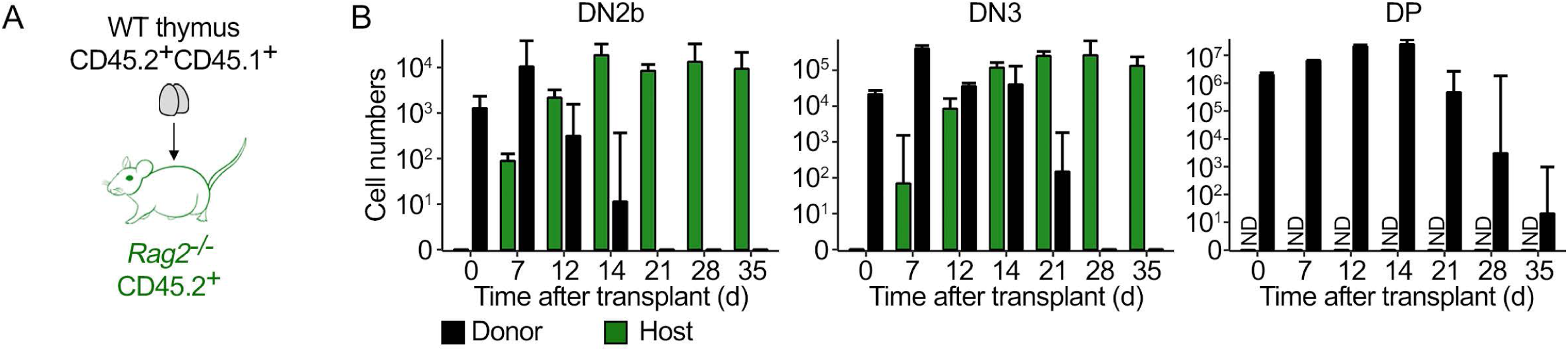
Kinetics of wild type thymopoiesis in *Rag2*^*-/-*^ recipients. **(A)** Schematics of the experimental design. **(B)** Thymus grafts were analyzed by flow cytometry at the indicated time points after transplant (in days, d) and the absolute number of DN2b (left), DN3 (middle) and double positive (DP, right) were determined at the indicated time points after transplant (in days, d). Donor and host derived thymocytes are represented in black or green bars, respectively. ND = not detectable. Data were pooled from several experiments using a total of 4-6 grafts per time point. Shown is the median + 95% confidence interval.

### Cell competition occurs at DN2 and DN3 stages of differentiation

Although donor and host ETP and DN2a did not coexist at the analyzed time points (Fig. 1), it was still possible that host cells at those developmental stages could be involved in the inhibition of thymus autonomy by competing with donor cells further downstream in differentiation. To test that hypothesis we compared thymopoiesis in wild type thymi transplanted into *Ccr7*^*-/-*^*Ccr9*^*-/-*^ recipients, which have a defect in thymus homing, and therefore a dramatically reduced ETP population (Liu et al., 2006; Zlotoff et al., 2010). Despite the defect in thymus homing, thymus autonomy was inhibited in these hosts (Fig. 3A). Consistent with the former results (Fig.1, 2), coexistence of donor and host thymocytes at DN2b and DN3 occurred before the most immature populations were reestablished by thymocytes of host origin (Fig. 3B). By 28 days post transplantation, the defect in ETP remained but the subsequent populations recovered, suggesting that a homeostatic equilibrium was sought (Fig. 3C). This may not be sustained in the long term, as the thymus of *Ccr7*^*-/-*^*Ccr9*^*-/-*^ mice has reduced numbers of DN cells (Zlotoff et al., 2010) (and data not shown). Nevertheless, these data demonstrate that ETP are unlikely to be involved in cell competition.

**Figure 3.**
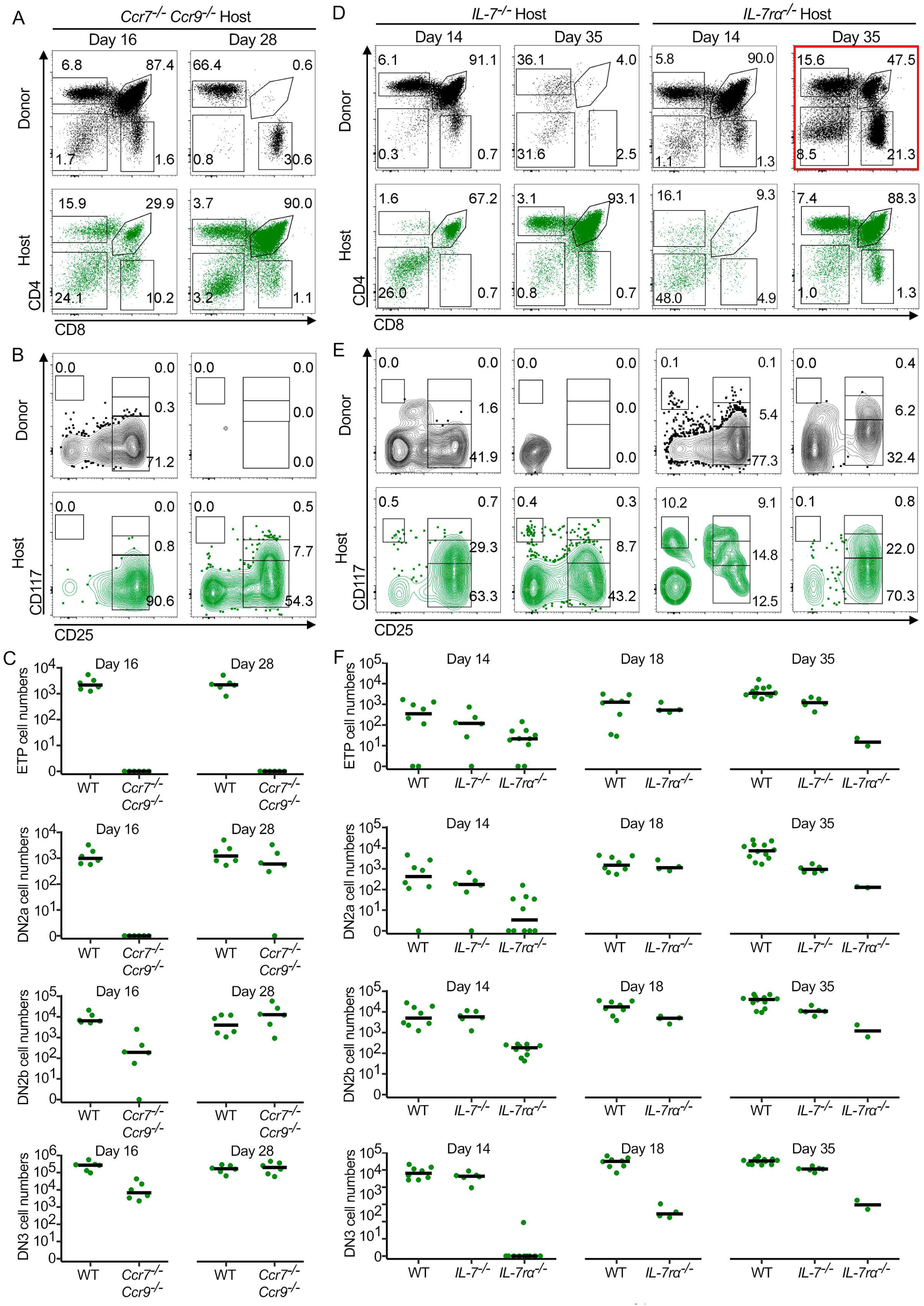
Cell competition is restricted to DN2 and DN3 via an IL-7 mediated response. **(A)** Representative CD4/CD8 profile of wild type thymi grafted into *Ccr7*^*-/-*^ *Ccr9*^*-/-*^recipients. Wild type donor (WT, black) and *Ccr7*^*-/-*^ *Ccr9*^*-/-*^ host (green) thymocytes are shown 16 and 28 days after transplant. **(B)** Representative CD117/CD25 profile of the thymus grafts in (A) pre-gated for negative CD4/CD8/lineage. **(C)** Absolute number of wild type (WT) or *Ccr7*^*-/-*^ *Ccr9*^*-/-*^ host derived ETP, DN2a, DN2b and DN3. **(D)** Representative CD4/CD8 profile of wild type thymi grafted into *IL-7*^*-/-*^ or *IL-7rα*^*-/-*^ recipients as indicated. Wild type donor (WT, black) and host (green) thymocytes are shown 14 and 35 days after transplant**. (E)** Representative CD117/CD25 profile of the thymus grafts in (D) pre-gated for negative CD4/CD8/lineage at the indicated time points. **(F)** Absolute number of host derived thymocytes in wild type (WT), *IL-7*^*-/-*^ and *IL-7rα*^*-/-*^ recipients at the indicated time points.

Response to IL-7 is essential for cell competition and the inhibition of thymus autonomy (Ballesteros-Arias et al., 2019; Martins et al., 2014). We sought to understand how IL-7 influences cell competition. For that purpose, we transplanted wild type thymi into *IL-7*^*-/-*^ versus *IL-7rα*^*-/-*^ mice (Fig. 3D). In these experimental settings, the hematopoietic progenitors seeding the thymus have been generated in the bone marrow in the absence of IL-7/IL-7r signaling (Peschon et al., 1994; von Freeden-Jeffry et al., 1995). After colonization of a wild type thymus, the deficient cells for the cytokine, *IL-7*^*-/-*^, can respond to IL-7 but cells deficient for the receptor, *IL-7rα*^*-/-*^, cannot. Interestingly, *IL-7*^*-/-*^ host thymopoiesis replaced that of donor origin by 35 days after transplantation, while that of *IL-7rα*^*-/-*^ host did not (Fig. 3D). The latter were indeed unable to prevent thymus autonomy, evidenced by the persistence of CD4+CD8+ double positive donor thymocytes 35 days after transplant (Fig. 3D, marked in red). Host thymopoiesis could be assessed at days 14 and 35 post transplantation, with significant differences between *IL-7*^*-/-*^ and *IL-7rα*^*-/-*^ at the early stages of differentiation (Fig. 3E). Although the numbers of ETP at day 14 differ between host *IL-7*^*-/-*^ and *IL-7rα*^*-/-*^, the most dramatic difference occurs for DN2 and DN3. At day 18, host *IL-7rα*^*-/-*^ thymocytes recover slightly but they were always reduced in comparison with the wild type counterparts (Fig. 3F). These data suggest that the numbers of the DN2 and DN3 populations are critical in the outcome of cell competition, and that it is unlikely that ETP are directly involved in this interaction. In other words, young DN2 and DN3 thymocytes of host origin replace the old thymocytes of donor origin that would otherwise be able to persist in an autonomous state. IL-7 is fundamental in cell competition potentially by defining the cell number of the populations involved. It remains to be tested whether the cell numbers per se define the outcome of the interaction or rather determine the availability of some limiting factor(s).

### DN2b and DN3 thymocytes adjust cell numbers transiently according to feedback from more mature developmental stages

Thymopoiesis is generally considered from the perspective that developing thymocytes follow a rather complex but predefined program. The prediction is that thymopoiesis and the kinetics of thymus turnover are expected to be constant. Much in line with these aspects is the directionality of cell competition that promotes thymocyte replenishment and thymus turnover. We sought to test whether thymocytes in general, and those involved in cell competition in particular, were capable of integrating information derived from the differentiation of more mature stages. To that end, we selected *Bcl2* transgenic mice that overexpress a human *BCL2* transgene in the T lymphocyte lineage (Katsumata et al., 1992; Strasser et al., 1991). *BCL2* overexpression delays apoptosis in thymocytes, mostly during positive selection (Akashi et al., 1997; Kondo et al., 1997), and we predicted that the kinetics of thymus turnover would be prolonged in time, as these thymocytes proliferate less than the non-transgenic counterparts (Mazel et al., 1996; O’Reilly et al., 1997). Thymi are similar between *Bcl2* transgenic and non-transgenic littermates (Figure S1A), with no changes in endogenous *Bcl2* expression (Figure S1B), and confirmed expression of the transgenic BCL2 in thymocytes (Figure S1C). This did not grossly affect thymus cellularity, although total number of ETP and CD8 single positive cells mildly increased (Figure S1D), and DN2b and DN3 slightly reduced when compared to wild type littermates (Figure S1D). No significant differences were detected in Annexin V-positive cells (Figure S1E). Instead, we detected a reduction in the percentage of proliferating *Bcl2* transgenic thymocytes (Figure S1F), in accordance with previous reports (Mazel et al., 1996; O’Reilly et al., 1997).

To test whether thymocyte subpopulations could detect and react to changes in the speed of differentiation of more mature populations, we compared thymus turnover in wild type versus *Bcl2* transgenic donor thymi following transplantation into wild type recipients (Fig. 4A). In this experimental setting, host derived thymocytes were always wild type and the difference was in the genotype of the donor thymus (Fig. 4A). The prediction was that if the kinetics of thymus seeding and early thymopoiesis would rely solely on the wild type genotype of the incoming cells, then host thymocytes would seed the thymus graft and differentiate identically in both conditions. Surprisingly, we observed that when wild type thymocytes differentiated in a *Bcl2* transgenic thymus, they did so with delayed kinetics as compared to that of wild type thymocytes in a wild type thymus graft (Fig. 4B, C). The delay in thymocyte replacement was most obvious for double positive thymocytes, seen in the reduced percentage of CD4+CD8+ double positive thymocytes measured between conditions (Fig. 4B). Furthermore, the curves describing donor and host derived double positive cells intersect at day 17 in the wild type, or 21 in the *Bcl2* transgenic grafts (Fig. 4C). Nevertheless, measuring thymocyte populations only by percentages could be misleading, as they could reflect the accumulation of the *Bcl2* transgenic donor rather than a difference in host derived thymocytes. Therefore, we quantified the absolute number of cells within each compartment over time and detected that wild type thymocytes of host origin were indeed transiently reduced in numbers at the DN2b and DN3 stages specifically in *Bcl2* transgenic thymus, as compared to their counterparts differentiating in wild type thymus donors (Fig. 4D). The delay in expansion of host derived DN2b and DN3 thymocytes correlated with the prolonged time that double positive *Bcl2* transgenic donor cells took to differentiate and progress out of this developmental stage (Fig. 4D). Consistent with the rescue from failed positive selection induced by the transgene (Strasser et al., 1991), donor double positive transgenic thymocytes indeed showed a slower kinetics of differentiation (Fig. 4D, grey bars). These experiments gave similar results to experiments of thymus transplants into *Bcl2* transgenic recipients (Figure S2A). Specifically, the kinetics of turnover in a *Bcl2* transgenic thymus was also delayed as compared to that in a wild type thymus graft in *Bcl2* transgenic hosts (Figure S2B, C), and that was reflected in absolute cell number in each compartment over time (Figure S2D). In both sets of experiments, host derived thymocytes at the DN2b and DN3 stages were transiently reduced in numbers when differentiating in the *Bcl2* transgenic thymus (Fig. 4, Figure S2). Altogether, our results indicate that DN2b and DN3, the majority of cells that are involved in cell competition, adjust their numbers transiently, according to the speed of differentiation of more differentiated thymocytes, presumably the CD4+CD8+ double positive. This is a general property that does not depend exclusively on the genotype of the recipient bone marrow derived cells, but rather reflects their response to the environment that they encounter in the thymus.

**Figure 4.**
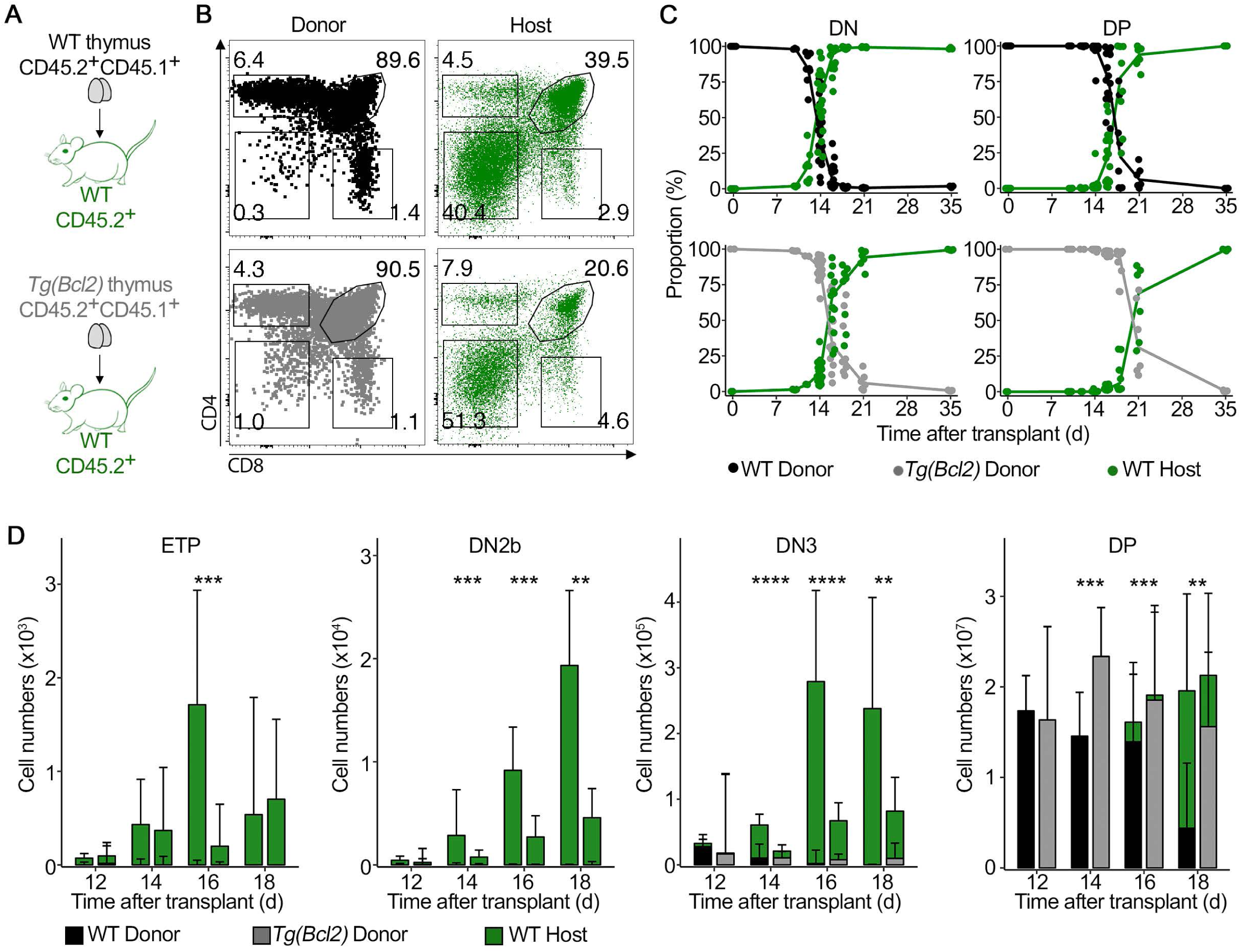
Wild type thymocytes have a slower kinetics of thymus turnover when seeding a *Bcl2* transgenic thymus. **(A)** Schematics of the experimental design. **(B)** Representative FACS plots from thymus grafts 21 days after transplant. Top left: donor wild type thymocytes; bottom left: donor *Tg(Bcl2)* thymocytes; top right: host wild type thymocytes in wild type thymus graft; bottom right: host WT thymocytes in *Tg(Bcl2)* thymus graft. **(C)** Proportion of donor and host thymocytes in double negative (DN) and double positive (DP) thymocytes in non-transgenic (top row) versus *Tg(Bcl2)* grafts (bottom row). **(D)** Absolute number of host derived wild type (green bars) and donor (wild type in black, *Tg(Bcl2)* in grey) thymocytes in the indicated populations and time points after transplant. For each time point, non-transgenic grafts are on the left, and *Tg(Bcl2)* are on the right. (D) Wilcoxon signed-rank test. *p≤0.05, **p≤0.01, ***p≤0.001, **** p<0.0001. Data were pooled from several experiments using a total of 4-10 grafts per time point. Shown is the median + 95% confidence interval. See Figure S1 for characterization of *Bcl2* transgenic thymi. See Figure S2 for the kinetics of *Bcl2* transgenic thymopoiesis when seeding wild type thymus grafts.

In addition to the adjustment in cell numbers that occurred at DN2b and DN3 we also detected that host derived ETPs were increased in the wild type thymus grafts as compared to the same population in *Bcl2* transgenic thymus grafts. This was the case both in wild type (Fig. 4D) and *Bcl2* transgenic recipients (Figure S2D). Changes in the number of ETP could be explained either by differences in thymus seeding by bone marrow derived progenitors, or by differential expansion of the population of thymus seeding progenitors in the wild type thymus versus the transgenic thymus. We consider the latter explanation more likely, as the hosts are identical and only the thymus grafts differ.

### DN2b control cellularity by adjusting cell cycle duration, thereby determining the kinetics of thymus turnover

The observed differences in the absolute number of DN2b and DN3 could result from either differential cell death or proliferation rates. We measured apoptosis by Annexin V staining and detected a reduction in the percentage of Annexin V positive DN2 and DN3 wild type thymocytes in the *Bcl2* transgenic thymus grafts as compared to the same subsets differentiating in the wild type grafts (Fig. 5A). Next we assessed proliferation *in vivo* by EdU incorporation. By 14 days after transplantation, the earliest time point at which the differences in absolute number of DN2 and DN3 could be detected, host derived wild type DN2 thymocytes differentiating in *Bcl2* transgenic thymus grafts had a lower percentage of EdU-positive cells than DN2 differentiating in wild type thymus grafts (Fig. 5B). This difference was transient, as two days later the rate of proliferation at the DN2 increased and even surpassed that of cells differentiating in wild type thymi (Fig. 5B). To address whether the lower percentage of EdU incorporation could be explained by changes in the duration of cell cycle, we performed an EdU/BrdU double pulse assay (Fig. 5C). This sets a labeling pattern (Fig. 5D) that has been used for other cell types (Akinduro et al., 2018) and provides information about cell cycle dynamics (Fig. 5E). Focusing specifically in the DN2 cells, we used these data to apply a mathematical model that estimates the duration of cell cycle based on the dynamics of double pulse cell labeling (Methods section). The model consists of a two-state system in which cells can either be in S phase or outside S phase (Fig. 5E). The population outside S phase is composed of cycling cells that can be in G1, G2 or M. The model fits four values corresponding to the proportion of cells in each quadrant of the EdU/BrdU profile as a function of cell cycle length and the elapsed time between the two pulses. We could determine that DN2 prolonged cell cycle duration transiently in *Bcl2* transgenic thymi (Fig. 5F). Taken together, the data show that DN2b react differently to the thymus in which they differentiate mostly by altering the duration of cell cycle.

**Figure 5.**
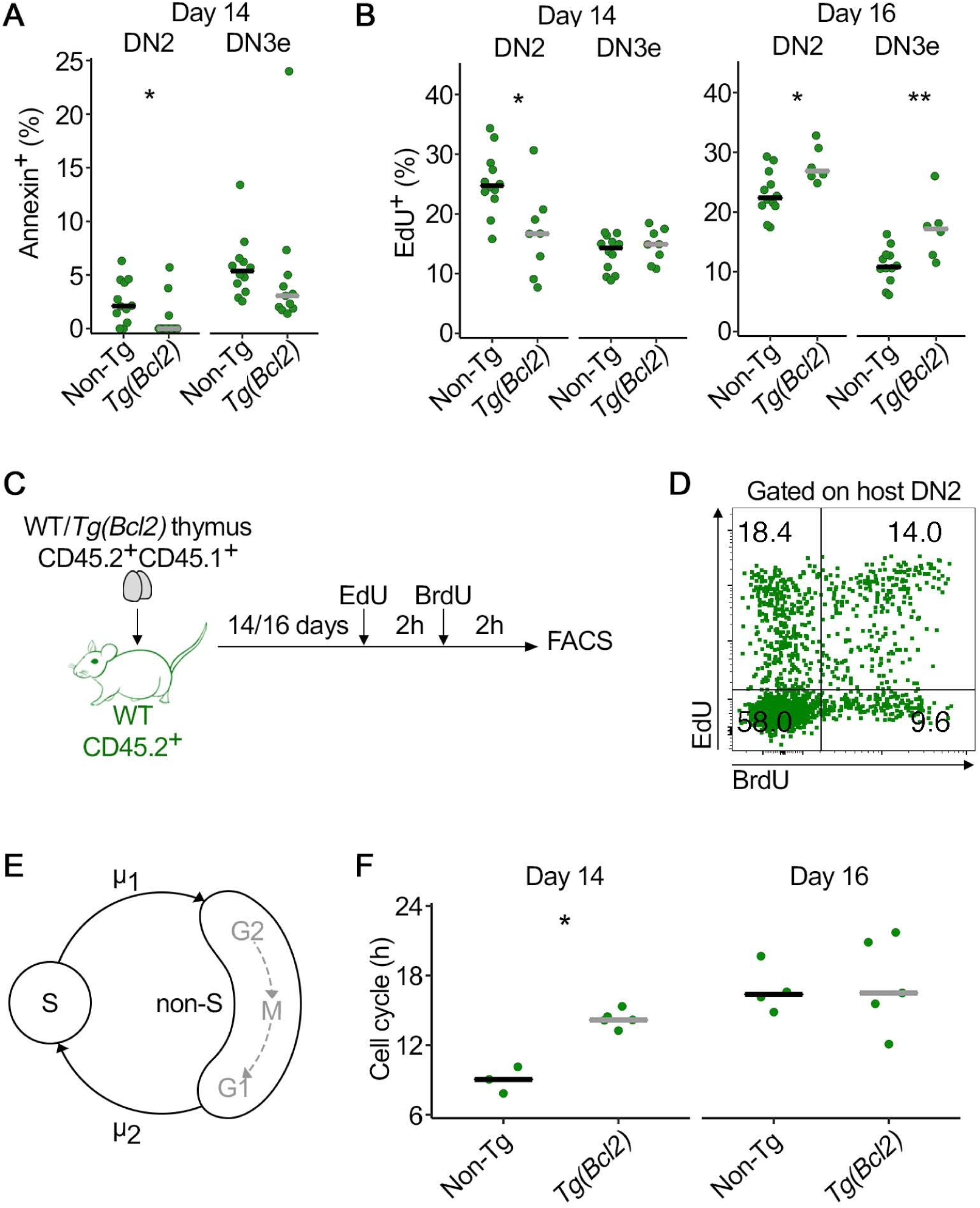
Thymopoiesis is regulated by proliferation and cell cycle duration of DN2b. Experimental design as in Fig. 4A and grafts were analyzed for **(A)** Annexin V 14 days after transplant. Data is from three independent experiments. **(B)** Proliferation was determined by EdU incorporation on days 14 and 16 after transplant, as indicated. Data is from three independent experiments. **(C)** Experimental design for the experiments in D) to F). Thymus transplants were performed as in Fig. 4A and at days 14 or 16 after transplant mice were injected i.p. with EdU, followed by BrdU after a 2-hour interval. Thymus grafts were analyzed two hours later. **(D)** Representative dot plot of EdU/BrdU profile of host DN2 thymocytes 14 days after transplant. **(E)** Schematics of the mathematical model used to calculate the duration of cell cycle. **(F)** Total cell cycle duration in hours (h) of wild type DN2 cells that differentiated in either non-transgenic (non-Tg) or *Tg(Bcl2)*, thymus grafts. Depicted are the results for day 14 (left panel) and 16 (right panel) after transplant. Each dot corresponds to the value for one graft and lines are medians. Wilcoxon signed-rank test. *p≤0.05, **p ≤0.01, ***p≤0.001, **** p<0.0001.

### The changes in proliferation and numbers of DN2b and DN3 depend directly on the presence of double positive thymocytes

The data so far was consistent with the idea that the transient response of the DN2b – slowing down cell cycle, with the resulting impact on cell numbers, thereby prolonging the kinetics of thymus turnover in the *Bcl2* transgenic thymus – was caused by the longer endurance of thymocytes at the double positive stage. It is reasonable to consider that double negative and double positive thymocytes could interfere with each other, since they both locate in the thymic cortex. To test this, we performed thymus transplantation experiments similar to those depicted in Figure 4 but in *Rag*-deficient background (Fig. 6A). *Rag2*^*-/-*^ thymocytes fail to differentiate beyond the DN3e stage, because they cannot rearrange the T cell receptor *loci*, so double positive thymocytes are never generated. The hematopoietic progenitors seeding the thymus grafts were *Rag2*^*-/-*^ and they seeded a *Rag2*^*-/-*^ thymus that was either *Bcl2* transgenic, or non-transgenic (Fig. 6A). In this experimental setting, thymopoiesis up to the DN3 stage progressed in both conditions (Fig. 6B), but the cell numbers of host derived DN2b and DN3 in *Bcl*2 transgenic or non-transgenic thymus grafts no longer differed significantly in the *Rag*-deficient background (Fig. 6C). This indicates that double positive thymocytes interfere with the pace of differentiation of the more immature thymocytes. Furthermore, no difference was detected in the percentage of proliferating DN2b and DN3 thymocytes (Fig. 6D).

**Figure 6.**
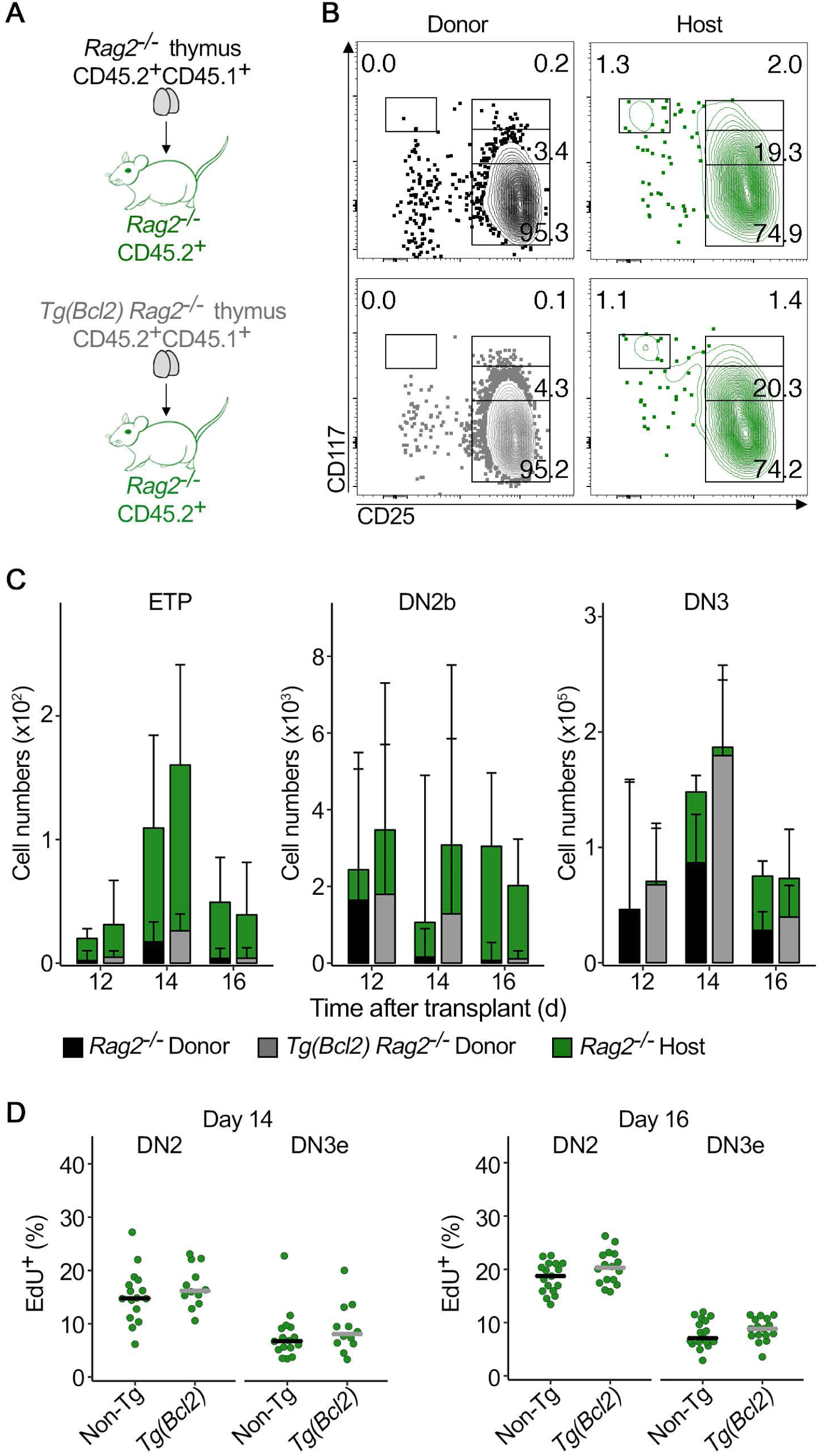
Double positive thymocytes interfere with the proliferation of DN2b and DN3. **(A)** Schematics of the experimental design. **(B)** Representative contour plots of CD117/CD25 profile of donor (left) and host derived (right) thymocytes in non-transgenic (top) or *Bcl2* transgenic (bottom) grafts 14 days after transplant. **(C)** Quantification of the total number of host *Rag2*^*-/-*^ (green bars) and donor (*Rag2*^*-/-*^ in black, *Rag2*^*-/-*^ *Tg(Bcl2)* in grey) thymocytes at the indicated time points after transplant. For each time point, *Rag2*^*-/-*^ grafts are on the left and *Rag2*^*-/-*^ *Tg(Bcl2)* are on the right. Data are median +95% confidence interval and at least 6 grafts per time point are depicted. **(D)** Proliferation was determined in a 2-hour pulse EdU incorporation assay at the indicated time points after transplant. Quantification is depicted for host derived, *Rag2*^*-/-*^ DN2 and DN3e thymocytes. (D) Points represent individual grafts and bars the medians. (C, D) Wilcoxon signed-rank test. *p≤0.05, **p≤0.01, ***p≤0.001, **** p<0.0001. Data are three independent experiments for day 14 and 16. Day 12 is from one experiment.

The fact that the cell numbers of host derived DN2b and DN3 thymocytes were the same in *Rag2*^*-/-*^ *Bcl*2 transgenic and *Rag2*^*-/-*^ non-transgenic thymus grafts suggests that double positive thymocytes influence the expansion of DN2b. The result of that interaction is determining for the pace of differentiation at the double negative stages, which in turn defines overall thymus turnover.

### The changes in proliferation and numbers of DN2b and DN3 are regulated by IL-7 availability

IL-7/IL-7r signaling is important for proliferation and survival of thymocytes at the DN2 stage, and leads to their differentiation into DN3 thymocytes (DiSanto et al., 1995; Peschon et al., 1994; von Freeden-Jeffry et al., 1995). Since IL-7 is limiting in the thymus (Barata et al., 2019), we reasoned that IL-7 availability could regulate proliferation, thereby controlling absolute cell numbers of DN2b and DN3 thymocytes. To test that hypothesis, we took advantage of IL-7r heterozygote (*IL-7rα*^*+/-*^) mice, that have normal thymopoiesis (Figure S3A) but express less surface IL-7r (Figure S3B). Since the levels of available IL-7 are controlled by cellular intake (Martin et al., 2017), *IL-7rα*^*+/-*^ thymi are predicted to have higher levels of IL-7. Consistently, sorted and cultured DN2-3 *IL-7rα*^*+/+*^ thymocytes consumed more IL-7 than *IL-7rα*^*+/-*^ (Figure S3C). We transplanted *IL-7rα*^*+/-*^ *Bcl2* transgenic versus *IL-7rα*^*+/-*^ (non-transgenic) thymi into wild type recipients (Fig. 7A). Thymocyte subpopulations of *IL-7rα*^*+/-*^ *Bcl2* transgenic versus *IL-7rα*^*+/-*^ (non-transgenic) littermates were very similar (Figure S3D). In this experimental setting, recipient derived thymocytes were always wild type and seeded *IL-7rα*^*+/-*^ thymus grafts, in which IL-7 was less limiting. If increased availability of IL-7 increases the differential proliferation of DN2b when differentiating in a *Bcl2* transgenic versus a non-transgenic thymus, then the prediction is that the differences detected previously (Fig. 4, 5) would subside. Indeed, no differences in cell numbers (Fig. 7B), percentage of proliferating cells (Fig.7C), and cell cycle duration (Fig.7D) were detected between wild type thymocytes in transgenic versus non-transgenic thymi. Hence, IL-7 availability regulates the differential proliferation of DN2b that occurs in response to double positive thymocytes.

**Figure 7.**
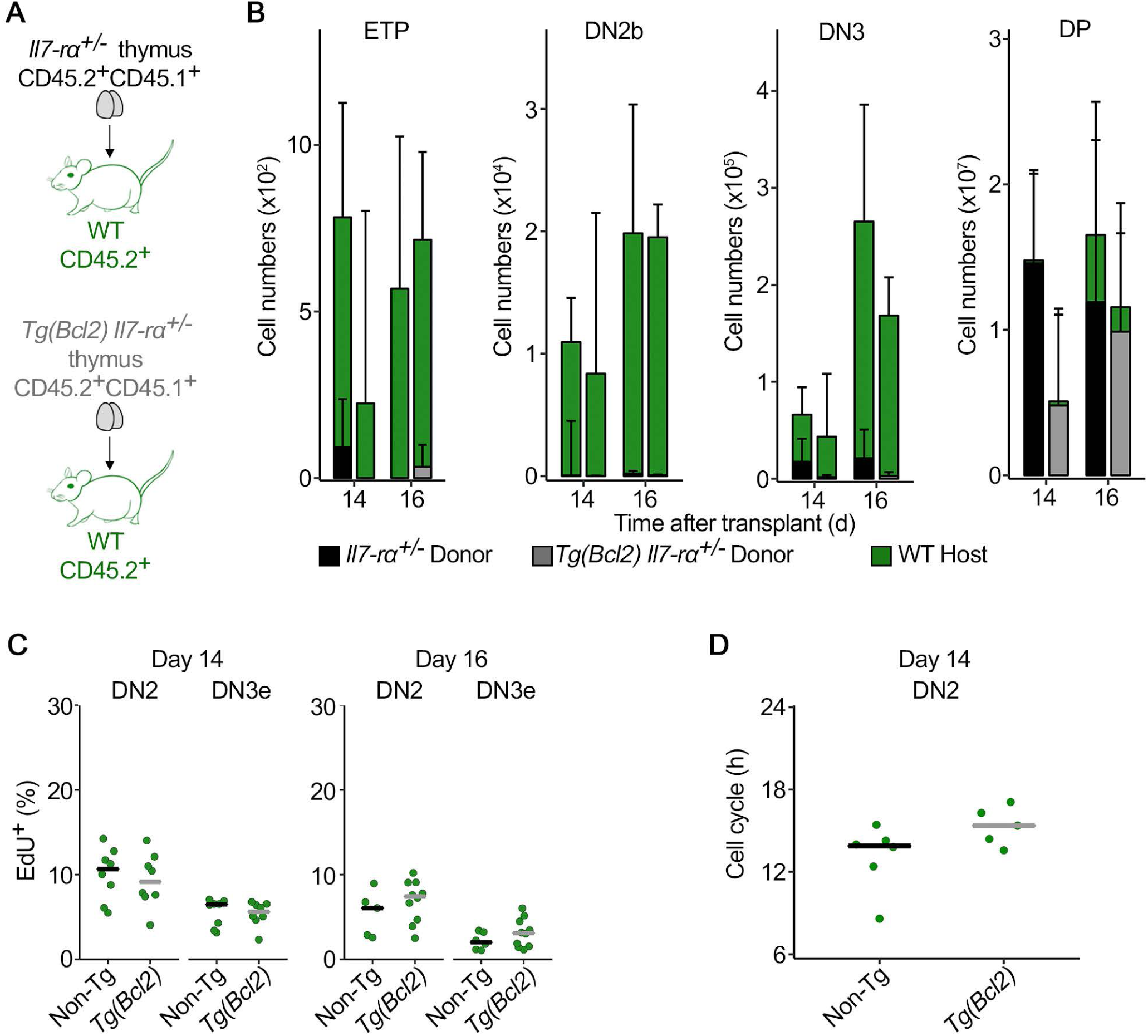
IL-7 availability in *IL-7rα*^*+/-*^ thymus grafts normalized observed differences in proliferation of DN2b. **(A)** Schematics of the experimental design. **(B)** Absolute number of host wild type (green) and donor derived (*IL-7rα*^*+/-*^, black, *IL-7rα*^*+/-*^ *Tg(Bcl2)*, grey) thymocytes at the indicated stages and time points. For each time point, *IL-7rα*^*+/-*^ grafts are on the left and *IL-7rα*^*+/-*^ *Tg(Bcl2)* are on the right. **(C)** Proliferation was determined in a 2-hour pulse EdU. Quantification is depicted for host derived, wild type DN2 and DN3e thymocytes at days 14 or 16 after transplant **(D)** Cell cycle duration for wild type host derived thymocytes that differentiated in *IL-7rα*^*+/-*^ with or without the *Tg(Bcl2)*, as indicated. (B) Data are median +95% confidence interval and at least 6 grafts per time-point are depicted. (C, D, E) Points represent individual grafts and bars the medians. Wilcoxon signed-rank test. *p≤0.05, **p≤0.01, ***p≤0.001, **** p<0.0001. (B, C) Data are at least two independent experiments per time point. See Figure S3 for characterization of *IL-7rα*^*+/-*^ and *IL-7rα*^*+/-*^ *Tg(Bcl2)* thymi.

Taken together, our data show that thymocytes at the DN2b control the kinetics of thymus turnover, reacting to the speed at which more differentiated progeny progresses in thymopoiesis. DN2b thymocytes impact on thymus turnover by adjusting cell cycle duration in an IL-7r dependent way, which directly affects absolute cell numbers within the DN2b. This propagates onto the numbers of DN3 and then subsequently onto the following stages of differentiation. Differentiation occurs in parallel with cell competition, and the number of DN2-DN3e thymocytes above a certain threshold (defined by response to IL-7) is determining for cell competition. In other words, the fine-tuning of DN2b expansion also impacts onto the kinetics of cell competition and the inhibition of thymus autonomy.

## DISCUSSION

Previous work has demonstrated that cell competition between thymocytes promotes thymus cellular turnover and is required to prevent thymus autonomy and T-ALL (Martins et al., 2014). Furthermore, for all conditions in which thymus autonomy was detected, the long-term consequence was the emergence of T-ALL (Ballesteros-Arias et al., 2019). Here we show that cell competition takes place in the thymic cortex, identify the cell populations involved, and show that their expansion is regulated by a negative feedback loop mechanism imposed by more mature thymocytes. Cell competition involves DN2 and DN3e thymocytes, inhibits thymus autonomy and therefore the emergence of T-ALL. This is the interaction that inhibits the persistence of DN3 thymocytes, which have the potential to self-renew and differentiate (Paiva et al., 2020). It remains to be seen if this involves direct cell-cell contacts, but the presence of thymocytes with different dwell time in the same regions of the thymus is consistent with that hypothesis. There is a step of regulation of DN2b thymocyte expansion that results in the control of proliferation, and thereby differentiation, of the more immature cells that are engaged in cell competition. Changes in cellularity of DN2b propagate sequentially, meaning that DN2b are effectively the cells defining the pace at which thymopoiesis proceeds, and thereby the kinetics of thymus turnover. It will be of interest to determine whether this interaction results from competition for limiting resources or whether there is a limit imposed by availability of space in the cortex.

Competition for limiting resources, including cytokines, growth factors or available niches play a central role in the regulation of the immune system and in homeostasis in the steady state (Surh and Sprent, 2008). In the thymus, competition for stromal niches occurs at early stages of differentiation and defines the number of available niches that can be occupied by thymus seeding progenitors at any given time (Prockop and Petrie, 2004). What thymocytes compete for is still elusive, but limiting factors like Notch ligands and IL-7 are likely important. In this context, *Notch1*^*+/-*^ progenitors generate reduced thymocyte progeny in mixed bone marrow chimeras, suggestive that Notch ligands are limiting, and potentially drive competition in the thymus (Tan et al., 2005). This is consistent with the competitive advantage of thymocytes overexpressing a Lunatic Fringe transgene, which confers increased sensitivity to Notch1 (Visan et al., 2006). While the step at which cell competition takes place could be explained by the presence of new incoming cells, that respond to IL-7 and therefore displace those cells of donor origin, this effect cannot be explained by differential expression of IL-7r (not shown). It will be necessary to further detail the mechanism regulating this interaction, but it is clear that the cell numbers of the populations involved are important. As for the negative feedback loop controlling DN2b expansion by double positive thymocytes, it is unlikely that DN2b and double positive thymocytes compete directly for access to the same ligands, as their requirements for differentiation do not seem to overlap (Yui and Rothenberg, 2014). Instead, it is plausible that the longer differentiation time of double positive *Bcl2* transgenic thymocytes translates into longer time of residence in the cortex, thereby interfering with the access of the proportionally very few DN2b to ligands expressed by the thymic stroma. Indeed, there is on average a relation of 5000 double positive thymocytes for each DN2b, and the longer residence of double positive thymocytes could interfere with cell migration throughout the cortex. The fact that IL-7 availability regulates expansion of DN2b under these conditions is not completely unexpected, as DN2b depend specifically on this cytokine to proliferate and differentiate (Cao et al., 1995; DiSanto et al., 1995; Peschon et al., 1994).

Further studies will focus on understanding the factors enabling thymus autonomy, and how autonomy permits leukemia initiation (Paiva et al., 2018). In this context, it is interesting that work from two independent groups using bone marrow chimeras points out that lymphoid malignancies emerge in *common gamma c* (γ_*c*_) deficient hosts reconstituted with wild type bone marrow cells in limiting dosages (Ginn et al., 2017; Schiroli et al., 2017). In such conditions, it is possible that the efficiency of thymus seeding is compromised, with interspersed periods that are refractory to thymus colonization, and thymus autonomy is established. We consider likely that any condition that enables thymus autonomy for a prolonged time period is permissive to T-ALL (Paiva et al., 2018).

Finally, cell competition was originally described and studied as a homeostatic process contributing to tissue homogeneity and optimal organ function (Amoyel and Bach, 2014; Claveria and Torres, 2016). The process we describe in the thymus is reminiscent of this role, even though the molecular mechanisms involved are probably different, as they ought to be cell context dependent. Furthermore, cell competition has been shown to protect from cancer in the mouse intestine, by extrusion of transformed epithelial cells into the lumen (Kon et al., 2017). Nevertheless, cell competition has also been implicated in cancer by promoting an advantage to tumor cells as opposed to the cells in the normal tissue (Eichenlaub et al., 2016; Madan et al., 2019; Suijkerbuijk et al., 2016). It will be of interest to determine whether a similar process plays a role in the initiation and progression of T-ALL.

## ACKNOWLEDGEMENTS

This work was supported by the Instituto Gulbenkian de Ciência (IGC), Calouste Gulbenkian Foundation, and the Portuguese National Research Council (Fundação para a Ciência e Tecnologia [FCT] Grant PTDC/BIA-BID/30925/2017 to VCM). CVR and RAP are PhD students of the IGC Integrative Biology and Biomedicine (IBB) PhD Program and are supported by individual FCT PhD Fellowships refs. PD/BD/139190/2018 and PD/BD/114341/2016, respectively. This work had the support of the research infrastructures Congento LISBOA-01-0145-FEDER-022170 and PPBI-POCI-01-0145-FEDER-022122, both co-financed by FCT and Lisboa2020, under PORTUGAL2020 agreement (European Regional Development Fund). We thank T Boehm for the *Ccr7*^*-/-*^ *Ccr9*^*-/-*^ mice and A Cumano for the OP9-Dll4. We thank J Howard, HJ Fehling, G Morata and RS Paiva for critical reading of the manuscript. We acknowledge V Correia for technical support. We thank the team of Animal House Facility of IGC for the outstanding support to this work, and acknowledge the Advanced Imaging Facility, Antibody, Histopathology and Flow Cytometry Units of IGC in supporting this work.

## AUTHOR CONTRIBUTIONS

CV Ramos designed and performed most experiments, analyzed data, implemented the mathematical model for duration of cell cycle and wrote the manuscript, L Ballesteros-Arias and JG Silva performed experiments and analyzed data, RA Paiva performed experiments, M Nogueira performed immunohistology, E Gjini designed the mathematical model and supervised its application to the data, J Carneiro advised on the model implementation and interpretation, and VC Martins conceived the study, designed research, designed experiments, analyzed data and wrote the manuscript. All authors edited and contributed to the final version of the manuscript.

## STAR Methods

## RESOURCE AVAILABILITY

### Lead contact

Further information and resource requests should be directed to and will be fulfilled by the Lead Contact, Vera C. Martins

### Materials availability

This study did not generate new unique reagents.

### Data and code availability

Code implementing cell cycle duration model and corresponding parameter estimation is freely available at https://github.com/LDL-IGC/CellCycleModel

## EXPERIMENTAL MODEL AND SUBJECT DETAILS

### Mice

C57BL/6J (B6, CD45.2^*+*^) mice were bred and kept at the Instituto Gulbenkian de Ciência (IGC) in a colony that is frequently renewed with mice purchased from Charles River. *Rag2*^*-/-*^ (Shinkai et al., 1992) originally purchased from Taconic were also bred and kept at the IGC. B6.SJL-*Ptprc*^a^ Pep3^b^/BoyJ (CD45.1^+^) mice, JAX stock #002014, here termed B6/Ly5.1, *IL-7rα*^*-/-*^ (Peschon et al., 1994), JAX stock #002295, and B6.Cg-Tg(BCL2)25Wehi/J (JAX stock #002320, here termed *Tg(Bcl2)*) were purchased from The Jackson Laboratory. *IL-7*^*-/-*^ (von Freeden-Jeffry et al., 1995) were backcrossed to B6 (Carvalho et al., 2001) and kept at IGC. *Ccr7*^*-/-*^ *Ccr9*^*-/-*^ were kindly provided by T. Boehm after crossing the original single mutants (Calderon and Boehm, 2011) and kept at IGC. The original *Ccr7*^*-/-*^ (Ma et al., 1998) had been purchased from The Jackson Laboratory, and the *Ccr9*^*-/-*^ (Benz et al., 2004) were generated at the MPI Freiburg, Germany. All other strains used in this work were obtained by crosses between the original strains described here. All thymus analyses were performed in mice between the ages of 3 and 10 weeks. *Bcl2* transgenic mice were analyzed with non-transgenic littermate controls. Male and female mice (in similar proportions) subjected to thymus transplantation experiments were between 5 to 10 weeks of age. Thymi for thymus transplants were harvested from newborn donors in a F1(B6xB6/Ly5.1) genetic background, unless stated that B6/Ly5.1 were used instead, and including the described mutations. All mice were bred and kept in individually ventilated cages in the SPF area of the mouse facility of the Institute. All animal experiments were approved by the Ethics Committee of the IGC – Fundação Calouste Gulbenkian and the Direção Geral de Alimentação e Veterinária (DGAV).

## METHOD DETAILS

### Thymus transplants

Thymus transplants were performed as described previously (Martins et al., 2014; Martins et al., 2012). Briefly, newborn thymus lobes were physically separated and kept in cold PBS until engraftment. Recipient mice of both genders aged 5 to 10 weeks were anesthetized with Ketamine (100mg/kg) and Xylazine (16mg/kg) and each recipient received one thymus grafted under the kidney capsule, with individual lobes at opposite ends. Anesthetized animals were given a subcutaneous injection of saline solution and allowed to recover in a heated pad. In experiments involving *Bcl2* transgenic and non-transgenic thymus transplants, the transplanted thymi were from *Bcl2* transgenic and wild type littermate controls. Likewise, experiments in which *Bcl2* transgenic and non-transgenic recipient mice were used, the experimental groups consisted of littermates of both genders.

### Cell preparation and flow cytometry

Single-cell suspensions were prepared in PBS/10%FBS. Cell numbers were determined using a hemocytometer and dead cells excluded by Trypan Blue staining. Cells were blocked for 15 minutes with mouse IgG (112mg/ml, Jackson ImmunoResearch) and stained with an antibody solution appropriately diluted during 30 minutes. Whenever necessary, after two consecutive washes, the cells were subjected to additional staining steps. The antibodies and streptavidin, purchased from Biolegend, were as follows: BCL2 Alexa 488 (100), Bcl2 PE (BCL/10C4), BrdU Alexa 647 (3D4), CD3ε bio (145-2C11), CD3ε APC-Cy7 (145-2C11), CD3ε FITC (145-2C11), CD4 PE (GK1.5), CD4 PE-Cy7 (GK1.5), CD4 BV421 (GK1.5), CD4 BV605 (GK1.5), CD8a BV711 (53-6.7), CD8a bio (53-6.7), CD8a FITC (YTS169.4), CD8 PerCP-Cy5.5 (53-6.7), CD11b bio (M1/70), CD11c bio (N418), CD19 bio (6D5), CD19 PE-Cy7 (6D5), CD19 BV711 (6D5), CD25 BV421 (PC61), CD25 BV605 (PC61), CD44-PerCP-Cy5.5 (IM7), CD44 BV711 (IM7), CD45.1 APC (A20), CD45.1 BV421 (A20), CD45.1 PE-Cy7 (A20), CD45.2 FITC (104), CD45.2 PerCP-Cy5.5 (104), CD45.2 PE (104), CD45.2 PB (104), CD71 FITC (RI7217), CD98 PE (RL388), CD117 APC (2B8), CD117 APC-Cy7 (2B8), CD127 APC (A7R34), βTCR Alexa 647 (H57-597), βTCR BV711 (H57-597), βTCR FITC (H57-597), βTCR PE (H57-597), γδTCR PE (GL3), γδTCR bio (GL3), Gr-1 bio (RB6-8C5), NK1.1 bio (PK136), NK1.1 A647 (PK136), Ter119 bio (TER-119), streptavidin APC-Cy7, streptavidin BV605 and streptavidin BV785. In the thymus, DN thymocytes were defined as CD4-negative, CD8-negative and Lineage-negative (cocktail composed of CD3, CD11b, CD11c, Ter119, Gr1, NK1.1 and CD19). Dead cell exclusion was done using Sytox Blue (Molecular Probes), Zombie APC-Cy7 or Zombie Pacific Orange (Biolegend). Intracellular cell staining was performed by fixing the cells in 4% PFA for 15 minutes at room temperature, followed by intracellular staining using True-Nuclear Transcription Buffer Set (Biolegend) according to manufacturer instructions. Identification of apoptotic cells by Annexin V staining was performed according to manufacturer instructions (Biolegend, 640934). Quantification of proliferation using EdU incorporation was accomplished by IP injection of 0.5mg EdU (Thermo Fisher Scientific E10187). Organs were collected two hours later and the staining procedure was done according to manufacturers’ instructions using the Click-it Plus EdU Alexa Fluor 488 Flow Cytometry Assay kit (Molecular Probes). Quantification of cell cycle duration using an EdU/BrdU double pulse was performed by consecutive i.p. injections of EdU (0.5mg) and BrdU (1mg, Sigma-Aldrich B5002) two hours apart. Organs were collected and analyzed two hours after the second injection. Following normal staining procedure and fixation in 4% PFA, cells were permeabilized for 15 minutes and incubated for 1 hour at 37°C in *DNase I* (300ug/ml PBS, Sigma Aldrich). After washing the staining procedure for EdU was done according to the manufacturer’s instructions, followed by BrdU staining. All acquisitions were performed in a BD LSRFortessa X-20 cell analyzer using BD FACSDiva 8 software and analyses were done using FlowJo.

### Cell sorting

Thymocytes pooled from at least three *IL-7rα*^*+/+*^ or *IL-7rα*^*+/-*^ mice (aged 3 to 5 week-old) were incubated for 30 minutes with a biotinylated lineage-antibody cocktail: CD3ε bio (145-2C11), CD8a bio (53-6.7), CD11b bio (M1/70), CD11c bio (N418), CD19 bio (6D5), Gr-1 bio (RB6-8C5), NK1.1 bio (PK136) and Ter119 bio (TER-119). Cells were washed in PBS/10% FBS and incubated with Dynabeads Biotin Binder (ThermoFisher) for 40 minutes according to the manufacturer’s instructions. Lineage-depleted samples were then incubated with the following antibodies: CD4 PE-Cy7 (GK1.5), Streptavidin APC-Cy7, CD25 BV605 (PC61), CD44 PerCP-Cy5.5 (IM7), CD117 APC (2B8), CD71 FITC (RI7217), CD98 PE (RL388). Live/dead exclusion was made using Sytox Blue. Samples were sorted in a BD BACSAria II at a purity >99%. DN2 cells were gated as CD4^-^ Lineage^-^ CD25^+^ CD44^+/low^, DN3 early cells as CD4^-^ Lineage^-^ CD25^+^ CD44^-^ CD71^low/-^ CD98^low/-^, and DN3 late cells as CD4^-^ Lineage^-^ CD25^+^ CD44^-^ CD71^high/+^ CD98^high/+^

### Measurement of IL-7 consumption by ELISA

250 000 sorted DN2 and DN3 thymocytes from either *IL-7rα*^*+/+*^ or *IL-7rα*^*+/-*^ mice were cultured in 96 well plates for 24 hours onto OP9-Dll4 stromal cells with 100μl of supplemented IMDM medium containing 5ng/ml murine IL-7 and 5ng/ml murine Flt3-Ligand (Peprotech). The medium was stored at -80°C until analysis. ELISA for IL-7 was performed using Mouse IL-7 Quantikine ELISA Kit M700 (R&D Systems) according to manufacturer’s instructions.

### Immunofluorescence

Thymus grafts were embedded in OCT Compound and snap-frozen in liquid nitrogen. Tissue sections of 8μm were dehydrated in acetone and samples were preserved in a dry environment at -80°C until staining. Sections were incubated for 30 minutes with DAPI and mIgG in PBS/10%FBS at room temperature. Following two washes in PBS the samples were stained with CD25 PE (PC61), CD45.2 FITC (104) and Cytokeratin 8 A647 (1E8) overnight at 4°C. Antibodies were purchased from Biolegend. Stained sections were washed three times in PBS and mounted with Fluoromount-G (Thermo Fisher Scientific). Images were acquired on a commercial Nikon High Content Screening microscope, equipped with an Andor Zyla 4.2 sCMOS camera, using a 20x 0.75 NA objective.

### Image processing and quantification

Single images and thymus tiles were processed and stitched using Fiji (Schindelin et al., 2012). Donor and host CD25 positive cells were quantified using QuPath (Bankhead et al., 2017). Briefly, thymus sections were subdivided and categorized as cortical or medullary areas and capsule region. Damaged regions were excluded from analysis. Using a standardized threshold, the number of CD25-positive and CD45.2 positive cells was obtained and their (minimum) distance to the thymus capsule calculated.

### Estimation of cell cycle duration

The estimation of the cell cycle duration is based on a minimalist model of the cell cycle and labelling processes. A cell is assumed to be in one of two states corresponding to the cell cycle phases: it can be cycling through S or through G_2_/M/G_1_ phases. The cell transitions between S and non-S (G2/M/G1) states are described with the dynamics of a continuous time 2-state Markov Chain (Allen, 2010). The rate of exit from S phase is denoted by μ_1_, whereas the rate of entry into S phase is denoted μ_2_. The sojourn times in S and G2/M/G1 phases are random exponential variables with means respectively given by 1/μ_1_ and 1/μ_2_. The total duration of the cell cycle, denoted Δ is then obtained as: 

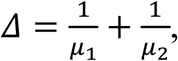

Under these conditions, the labelling dynamics of cells can be related to the time dependent transition probabilities between the two states of the Markov chain (S/non-S). For example, cells that take on both labels by the end of the experiment (++) are modelled as cells that were in S at time 0 (receiving the first label) and are found in S again at time t (receiving the second label). The proportions of cells in each quadrant (Fig. 5D) were thus rescaled, so that the probability of acquiring the second label is conditioned on the presence of the first DNA dye. Normalizing cell frequencies by their starting state (+ or -) for label 1, we have: *p*^+-^ is the proportion of EdU^+^BrdU^-^, *p*^++^=1-*p*^+-^ is the proportion of EdU^+^BrdU^+^, and *p*^-+^ is the proportion of EdU^-^BrdU^+^ and *p*^--^=1-*p*^-+^ is the proportion of EdU^-^BrdU^-^ cells. Taking into account the observed experimental frequencies after the initial labeling, the Markov chain model describes the proportion of labelled cells after time *t* (where *t* is the interval between the two labelling pulses) as follows:

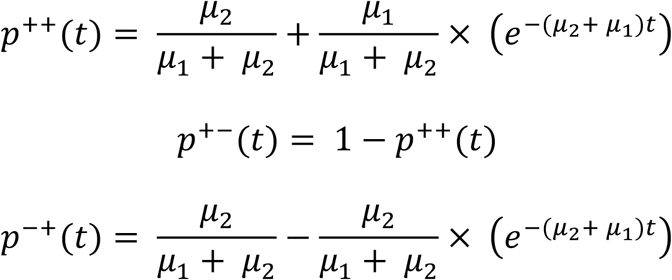

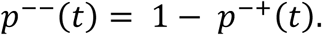

This model definition allows for an exact solution of the above equations, to find the unknown parameters (μ_1_, μ_2_) from the experimental proportions of labeled cells at t=2 hours. The model fitting (*μ*_1_*μ*_2_) was performed individually for each set of labeled cell proportions across different conditions. The estimates were gathered by the genotype of the graft and by day, and subsequently compared.

## QUANTIFICATION AND STATISTICAL ANALYSIS

Statistical analyzes were performed using R software. Statistical details of the experiments can be found in the figure legends. Significance values were computed using Wilcoxon rank-signed test. If data followed a normal distribution according to the Shapiro-Wilk test, unpaired two-tailed Student’s *t*-test was used. Prior to statistical analysis, thymus grafts that were found to be damaged or abnormal were excluded from the experimental groups.

