## Supplemental Figures and Figure Legends for "Cell competition, the kinetics of thymopoiesis and thymus cellularity are regulated by double negative 2 to 3 early thymocytes"

### Supplemental Figure 1

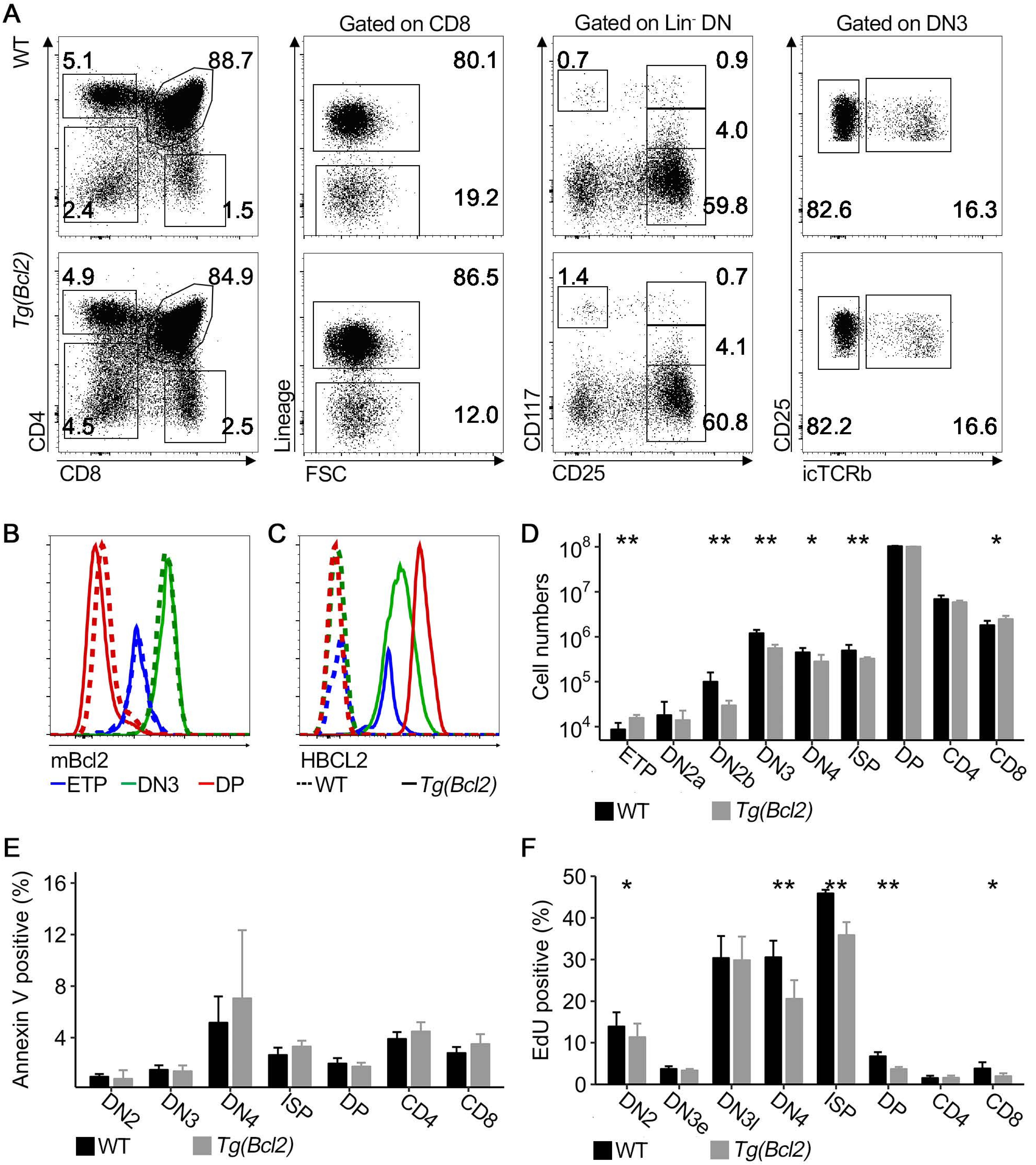


**Figure S1. Characterization of the *Bcl2* transgenic thymus,** Related to Figure 4. **(A)** The thymus of adult littermates that were wild type (WT, top row) or *Bcl2* transgenic thymus (bottom row) were analyzed by flow cytometry for the indicated markers, and **(B)** further analyzed for the expression of mouse Bcl2, and **(C)** human BCL2. **(D)** Absolute number of thymocytes was quantified in the indicated populations in *Bcl2* transgenic (grey bars) and in littermate wild type (WT) thymi (black bars). **(E)** Percentage of Annexin V positive thymocytes. **(F)** Percentage of EdU positive thymocytes at different developmental stages in adult *Bcl2* transgenic thymi (grey bars) and in littermate adult wild type (WT) thymi (black bars). EdU incorporation was assessed two hours after i.p. injection. (B, C, D, F) Data are representative of a minimum of two experiments. (D,F) Data are two independent experiments. (E) Data are one experiment. (D, E, F) Wilcoxon signed-rank test. *p≤0.05, **p≤0.01, ***p≤0.001, **** p<0.0001.

### Supplemental Figure 2

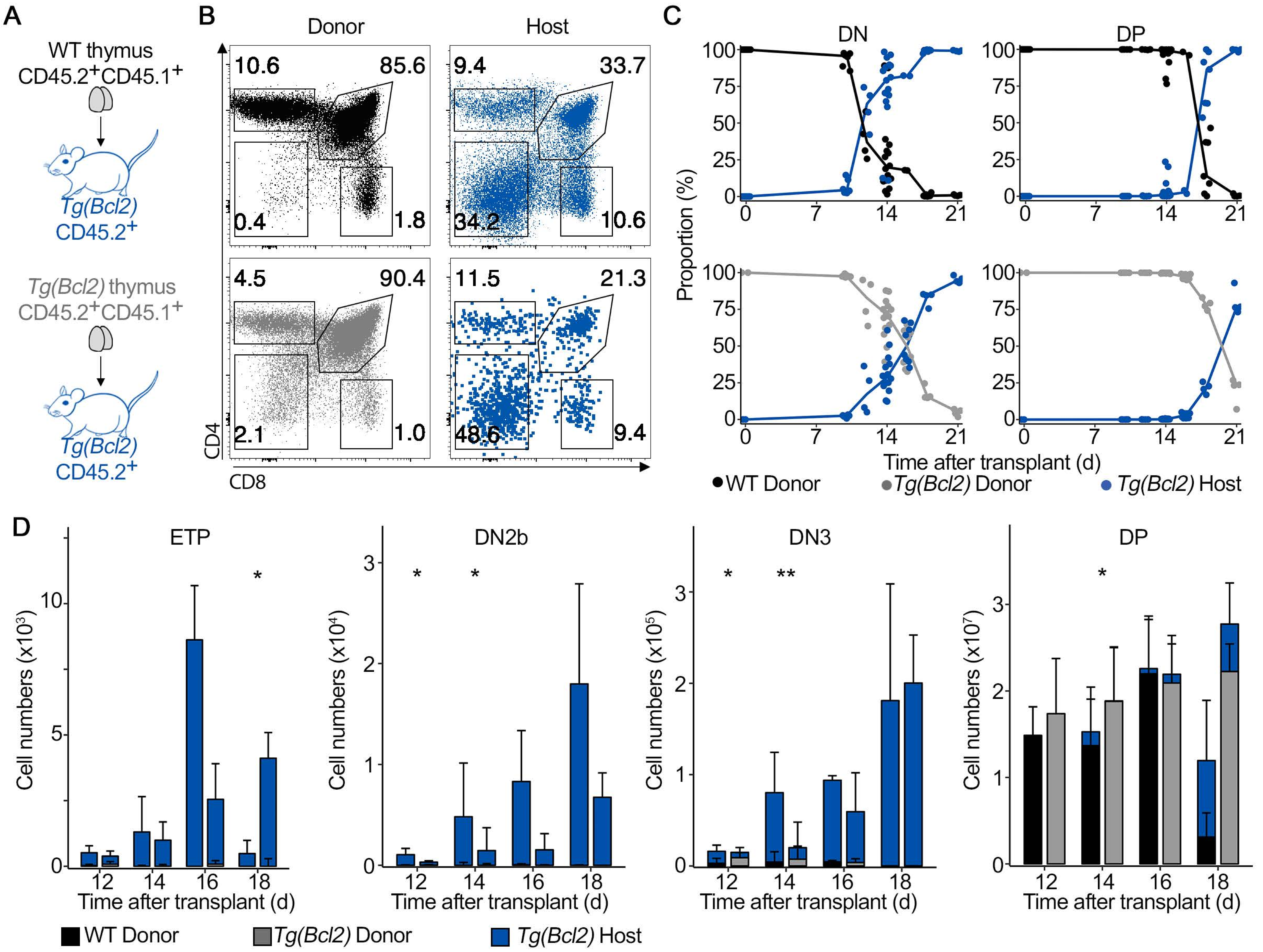


**Figure S2. Bcl2 transgenic thymocytes have slower kinetics of thymus turnover when seeding a *Bcl2* transgenic thymus,** Related to Figure 4. **(A)** Schematics of the experimental design. Non-transgenic or *Tg(Bcl2)* newborn donor thymus lobes were grafted under the kidney capsule of *Tg(Bcl2)* adult mice. One thymus was grafted per recipient. **(B)** Representative FACS plots of CD4/CD8 profile of live thymocytes from thymus grafts 14 days after transplant. Top left: donor wild type thymocytes; bottom left: donor *Tg(Bcl2)* thymocytes; top right: host *Tg(Bcl2)* thymocytes in donor wild type thymus; bottom right: host *Tg(Bcl2)* thymocytes in donor *Tg(Bcl2)* thymus. **(C)** Proportion of donor and *Tg(Bcl2)* host derived thymocytes in Lin- double negative (DN, left column) and CD4+CD8 double positive (DP, right column) during thymus turnover in wild type grafts (top row) and *Tg(Bcl2)* grafts (bottom row). **(D)** Absolute number of *Tg(Bcl2)* host (blue bars) and donor (wild type = black bars, *Tg(Bcl2)* = gray bars) thymocytes in different developmental compartments during thymus turnover. For each time-point, left bar-plots correspond to wild type grafts and right bar-plots correspond to *Tg(Bcl2)* grafts. (D) Wilcoxon signed-rank test. *p≤0.05, **p≤0.01, ***p≤0.001, **** p≤0.0001. Data were pooled from several experiments using a total of 4-6 grafts per time point, except for one group at day 16 with two datapoints. Shown is the median+95% confidence interval.

### Supplemental Figure 3

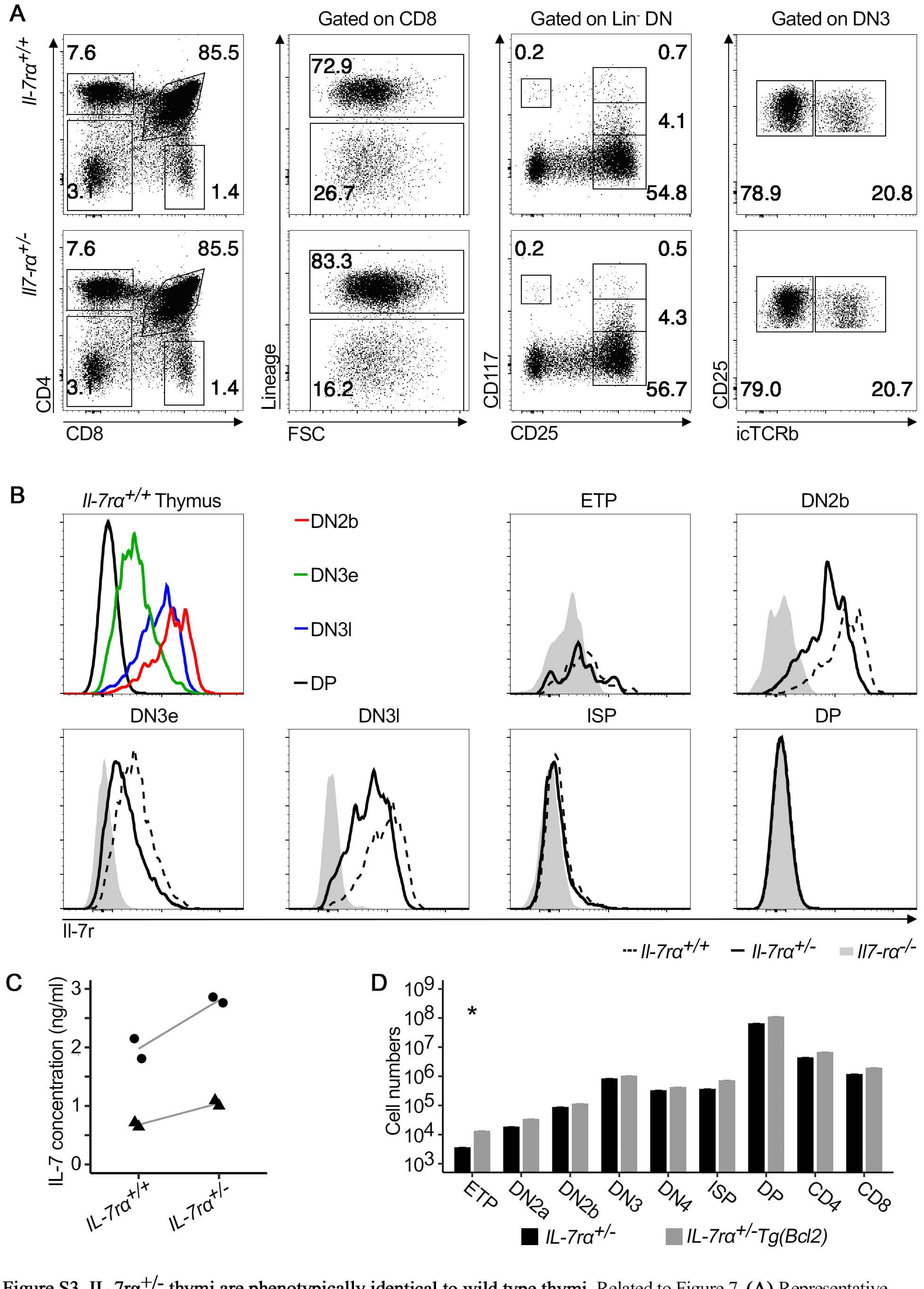


**Figure S3. IL-7rα$^{+/-}$ thymi are phenotypically identical to wild type thymi,** Related to Figure 7. **(A)** Representative FACS plots of thymocyte subpopulations in *IL-7rα$^{+/+}$* or *IL-7rα$^{+/-}$* thymi. **(B)** Histograms show Il-7rα expression in the indicated populations in an adult wild type (WT) thymus (left panel) and comparison of surface Il-7rα in different thymocyte subpopulations between *IL-7rα$^{+/-}$* (solid line), *IL-7rα$^{+/+}$* (dashed line) and *IL-7rα$^{-/-}$* thymi (grey histogram). **(C)** 250000 DN2 and DN3 thymocytes from IL-7rα$^{+/+}$ or IL-7rα$^{+/-}$ were sorted and co-cultured onto OP9-Dll4 with 5ng/ml murine IL-7. IL-7 concentration in the media was measured after culture for 24 hours. Different symbols correspond to two independent experiments. Lines connect the mean for each genotype. **(D)** Absolute number of thymocytes was quantified in the indicated populations in *IL-7rα$^{+/-}$* (black bars) and *IL-7rα$^{+/-}$ Tg(Bcl2)* (grey bars) littermate mice. Data are one experiment with three mice per group. Wilcoxon signed-rank test. *p≤0.05, **p≤0.01, ***p≤0.001, **** p≤0.0001.
